# Engineering clinically-approved drug gated CAR circuits

**DOI:** 10.1101/2020.12.14.419812

**Authors:** Hui-Shan Li, Nicole M. Wong, Elliot Tague, John T. Ngo, Ahmad S. Khalil, Wilson W. Wong

**Author notes:** Equal Contribution (Hui-Shan Li, Nicole M. Wong).

## Abstract

Chimeric antigen receptor (CAR) T cell immunotherapy has the potential to revolutionize cancer medicine. However, excessive CAR activation, lack of tumor-specific surface markers, and antigen escape have limited the safety and efficacy of CAR T cell therapy. A multi-antigen targeting CAR system that is regulated by safe, **clinically-approved** pharmaceutical agents is urgently needed, yet only a few simple systems have been developed, and even fewer have been evaluated for efficacy *in vivo*. Here, we present NASCAR (<u>N</u>S3 <u>AS</u>sociated CAR), a collection of inducible ON and OFF switch CAR circuits engineered with a NS3 protease domain deriving from the Hepatitis C Virus (HCV). We establish their ability to regulate CAR activity using multiple FDA-approved antiviral protease inhibitors, including grazoprevir (GZV), both *in vitro* and in a xenograft tumor model. In addition, we have engineered several dual-gated NASCAR circuits, consisting of an AND logic gate CAR, universal ON-OFF CAR, and a switchboard CAR. These engineered receptors enhance control over T cell activity and tumor-targeting specificity. Together, our comprehensive set of multiplex drug-gated CAR circuits represent a dynamic, tunable, and clinically-ready set of modules for enhancing the safety of CAR T cell therapy.

## INTRODUCTION

CAR-expressing T cells are a revolutionary FDA-approved cancer immunotherapy(Abramson et al., 2017; Brown et al., 2016; Davila et al., 2014; Maude et al., 2014; O’Rourke et al., 2017; Qasim et al., 2017; Rapoport et al., 2015). However, these engineered T cells can display adverse side effects such as cytokine release syndrome and cerebral edema, which can lead to fatalities(Brudno and Kochenderfer, 2016; Morgan et al., 2010). Current options for controlling adverse side effects involve systemic immunosuppression through drug administration (e.g., IL-6 inhibitor, sterols, or tyrosine kinase inhibitors)(Bonifant et al., 2016; Davila et al., 2014; Kumar et al., 2001; Le et al., 2018), or eradication of the engineered T cells through activation of a kill switch installed in the T cells(Di Stasi et al., 2011). Although these strategies can mitigate side effects, they adversely interfere with the therapy and native abilities to fight cancer, often causing patients to succumb to tumors after the intervention (Brudno and Kochenderfer, 2019). Furthermore, re-administration of the engineered T cells is undesirable and at times not feasible because of the prohibitive cost and complications involved in grafting T cells. Therefore, while a kill switch is useful as a last resort safety switch, it would be highly advantageous to reversibly regulate the activity of engineered T cells *in vivo* without sacrificing them when preventing or minimizing the onset of severe side effects.

In a similar principle to classic inducible gene switches, there are two types of controllable CARs - ON and OFF switches. Each type has its own unique advantages and therefore both are needed. ON switches are beneficial when off-target toxicity is an issue and T cells should only activate when dictated. However, in CARs deemed relatively safe, an OFF switch may be preferable. This would result in CAR T cells being active by default, but allows for temporary suppression of T cell activity or an immediate response to CRS if necessary. Furthermore, given that CRS can progress rapidly and a therapy can never be too safe, both ON and OFF switch capabilities in the same CAR would be beneficial for maximum controllability and safety.

Due to their recognized importance to CAR T cell development, small-molecule or protein inducible ON switch CARs (Juillerat et al., 2016; Kudo et al., 2014; Ma et al., 2016; Urbanska et al., 2012; Wu et al., 2015) and OFF switch CARs (Giordano-Attianese et al., 2020) have been developed. While these controllable CARs each have their unique advantages, undesirable properties of previously evaluated chemical inducers may limit their utility in clinical applications. Some of the systems developed use small molecules or proteins as inducers that have poor pharmacokinetics or tissue penetrance. Furthermore, almost all the existing systems use inducers that target human proteins and are not clinically-approved, and as such use drugs that have undetermined safety profiles. Additionally, current inducible system designs are limited in their versatility, restricting their implementation to either an ON or OFF switch, but not both. Other critical features, such as targeting multiple antigens to improve specificity or mitigate antigen escape, have yet to be incorporated into any drug inducible system. Therefore, an ideal approach to generate a clinically-relevant inducible CAR is to develop a system that uses an FDA-approved drug with favorable toxicity and pharmacokinetics profiles, and which can target multiple antigens.

To develop inducible CARs with safe and clinically-approved drugs, we leveraged the NS3 (Non-Structural Protein 3) system to create <u>N</u>S3 <u>AS</u>sociated CAR (NASCAR) systems. NS3 is a critical protein to the HCV life-cycle, which proteolytically cleaves the viral polyprotein at junction sites of non-structural proteins downstream of itself. Many regulatory agencies around the world, such as the US-FDA and Chinese FDA, have approved multiple drugs (e.g. grazoprevir (GZV) and danoprevir (DNV)) to inhibit the proteolytic activity of NS3(De Clercq and Li, 2016). Notably, GZV has an exceptional safety profile and is typically administered at 100 mg/day for 12 weeks, making it an ideal small molecule for clinical applications(Bell et al., 2016). This feature of drug-gated control over proteolytic activity has been exploited in other synthetic biology applications by generating drug-controllable imaging tags or degrons using the NS3 domain(Chung et al., 2015; Jacobs et al., 2018; Lin et al., 2008; Tague et al., 2018).

The computational design of ‘chemical reader’ systems, such as protein domains that are able to recognize liganded NS3 complexes bound to specific drugs, have also enabled the development of inducible dimerization and dissociation systems(Foight et al., 2019). Furthermore, using a peptide originally designed as an NS3 inhibitor(Kügler et al., 2012), along with a catalytically dead mutant of NS3, it is possible to form a heterodimerizing pair whose binding can be disrupted by GZV(Cunningham-Bryant et al., 2019). These protein engineering and chemical biology tools can be used to further increase the design and clinical potential of NASCAR systems.

Leveraging the unique advantages of the NS3 system, we have created a diverse set of controllable CARs with distinct functionalities. We have generated ON switch and OFF switch CARs, demonstrating both switch activities in a xenograft mouse tumor model. In addition, to improve tumor specificity and address antigen escape-caused relapses, we have developed several advanced dual-gated CAR circuits, which can either sense multiple antigens under AND-gate control or switch target antigen when desired. Combining our ON switch technology with a small molecule-controllable degradation system, we created a dual-gated AND CAR system that can enhance tumor targeting specificity by targeting two antigens. Furthermore, we have employed our Split, Universal and Programmable (SUPRA) CAR system(Cho et al., 2018) with the NASCAR OFF switch to create a universal ON-OFF switch CAR. The specificity of the SUPRA CAR is achieved through an adaptor protein, thus addressing the issue of antigen escape by retargeting tumors using alternative adaptor proteins. This is beneficial because the T cells would not need to be re-engineered. Our universal ON-OFF CAR has the added safety feature that it can be shut-off with GZV. Drug-gated control is particularly important for universal CARs because of the added risk that the additional target antigens are not sufficiently tumor-specific. Lastly, we have created a “switchboard” NASCAR system that is able to redirect T cell antigen specificity by inducibly recruiting the NS3 domain to different “reader” proteins through administration of either GZV or DNV. This dual-specific CAR system is genetically encoded into the T cells and does not require additional delivery of a protein adaptor, simplifying clinical administration though at the expense of reduced targeting flexibility. Here, we describe the efficacy of these various CAR circuits at regulating T cell activity, demonstrating how a drug-inhibited viral protease domain can extend and improve the versatility of CAR designs.

## RESULTS

### ON NASCAR

In an initial approach, we developed an ON switch CAR by inserting the NS3 cis-protease into the CAR framework (ON NASCAR) producing a single fusion protein in which the linkage between the scFv and signaling domains could be precisely controlled via drug-mediated NS3 inhibitor. In this design, the NS3 domain is flanked by NS5A/5B and NS4A/4B cleavage sites, and in the absence of a protease inhibitor (e.g. GZV), cis-proteolysis by the inserted enzyme is anticipated to produce a fragmented CAR, which we hypothesized would be unable to transduce any activation signals in response to targeted antigens. In the presence of NS3 inhibitor, however, we anticipated that blockage of protease activity would result in the formation of full-length and signaling-competent CARs, which would be able to mediate T cell activation following engagement with their specified antigen targets (**Figure 1A**). Using an anti-CD19 CAR as a model system, we generated three NS3-containing ON CAR constructs, varying the positioning of the NS3 domain within the receptor in order to identify optimal insertion configurations (**Figure S1A**).

**Figure 1:**
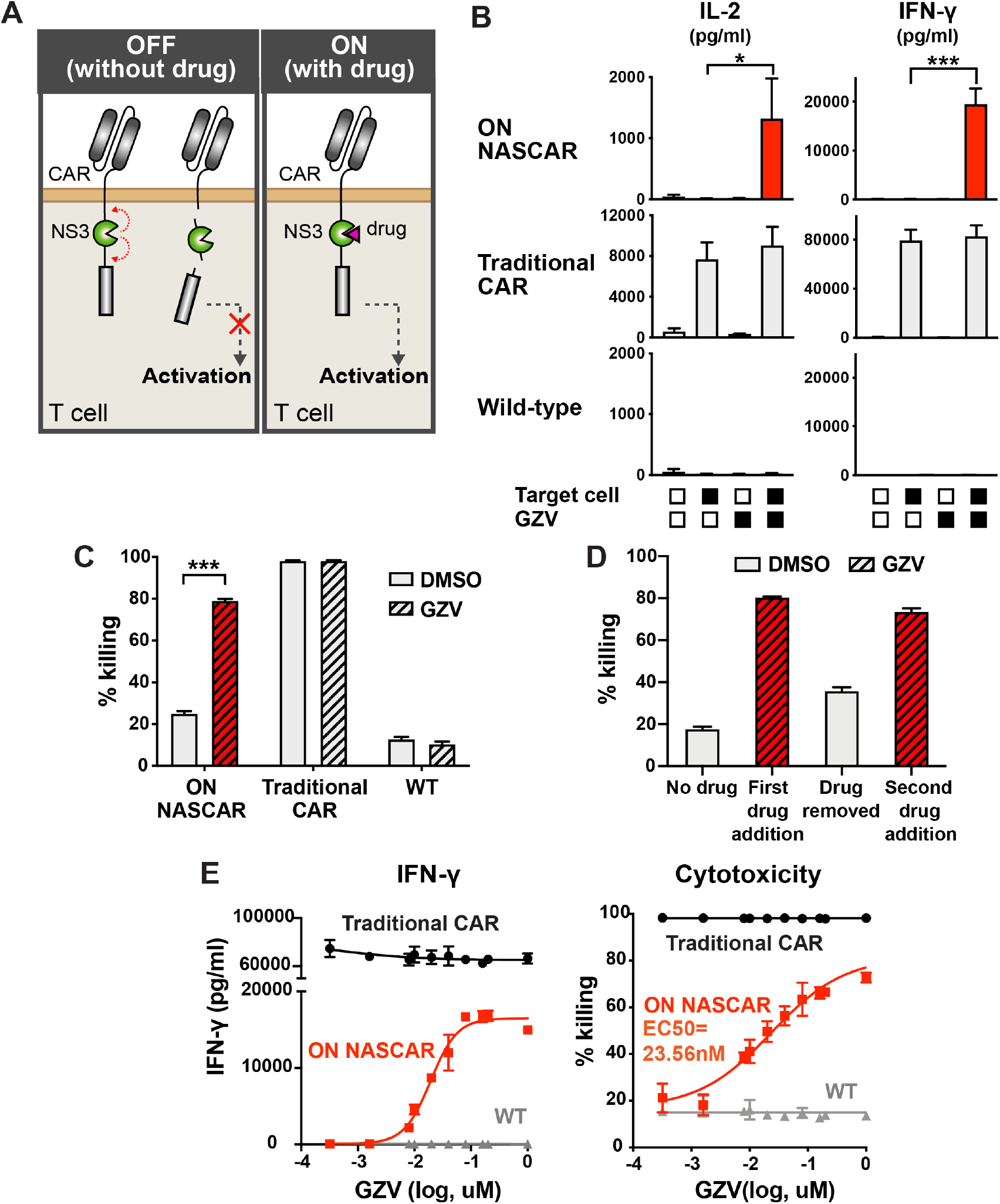
Characterization of the ON NASCAR. **A**) Schematic of the ON NASCAR mechanism. **B**) Primary T cell lines were treated with combinations of NS3 inhibitor and target NALM6 cells, and cytokine levels quantified (mean ± s.d., n = 3, \**P* < 0.05, \*\**P* < 0.01 and \*\*\**P* < 0.001). **C**) Comparison of cytotoxicity levels of the ON NASCAR with traditional CAR and wild-type cells (mean ± s.d., n = 3, \**P* < 0.05, \*\**P* < 0.01 and \*\*\**P* < 0.001). **D**) Target cell killing ability of ON NASCAR T cells when treated with drug, drug is washed out, and drug is reintroduced after 5 days (mean ± s.d., n = 3). **E**) GZV dose response profile of various cell lines, as measured in cytokine levels and cell killing (mean ± s.d., n = 3).

Next, we evaluated the signaling activities of the receptors. The three ON NASCAR versions were introduced into human primary T cells and activated with CD19+ NALM6 cells in the presence and absence of GZV. Activation of all three led to the increase of CD69 expression, cytokine production, and cytotoxicity (**Figure 1B, 1C, and S1C**). In comparison, a traditional CAR (with no NS3 domain) and wild-type T cells did not respond to GZV. Of the three configurations tested, the most effective version had the NS3 domain placed between the two costimulatory domains CD28 and 4-1BB. This design was therefore selected as the final ON CAR design moving forward (**Figure S1B**). The ON NASCAR can also be activated by three other FDA-approved NS3 inhibitors (glecaprevir, simeprevir and boceprevir)(McCauley and Rudd, 2016), with grazoprevir and glecaprevir able to induce cell killing at lower concentrations (**Figure S1E**). Based on the cytotoxicity dose response curve, the EC50 of the NS3 CAR to GZV was found to be 23.56 nM (**Figures 1E and S1D**). To determine the reusability of the ON NASCAR, we tested CAR reactivation after it had been switched on once already. We induced ON NASCAR T cells with GZV, confirming killing of CD19+ target cells, and then removed drug, which led to a decrease in cytotoxicity. When GZV was re-introduced, we found that killing efficiency increased to similar levels as before (**Figure 1D**), demonstrating the ability of the ON CAR to re-activate following previous stimulation. Interestingly, we also found that, while GZV is able to effectively regulate NASCAR activity, the level of cytokine release is lower than that observed with traditional (uninducible) CARs (**Figure 1B**). This is likely due to altered receptor expression of larger (NS3-containing) CAR constructs, the size of which is known to impact lentivirus packaging capability and consequent expression in cells (Kumar et al., 2001). However, since cytotoxicity levels are still comparable with traditional CARs, and exaggerated production of cytokines is the main cause of CRS, the tempered cytokine production could be considered a beneficial feature. Finally, we validated the generality of the ON NASCAR design by evaluating signaling and killing efficiency for a second scFv: an anti-Her2 scFv. The anti-Her2 ON NASCAR led to elevated production of CD69 and IL-2 in the presence of target cells and GZV (**Figure S2A**), and comparable killing efficiency and GZV dose-responsiveness as that of the anti-CD19 ON CAR (**Figures S2B and S2C**).

CARs are functional in other immune cell types and have immense therapeutic potential beyond cancer treatment. Therefore, it is also important to ensure that our ON NASCAR is functional in other cell types. Of particular interest is the usage of CARs in regulatory T (Treg) cells, where they can suppress immune responses. Treg cells are responsible for maintaining immune homeostasis and tolerance, and preventing autoimmunity. Due to their unique role in the immune system, Treg cells have been under investigation as therapeutic agents for treating autoimmune diseases, such as type 1 diabetes, or preventing organ transplant rejection(Kohm et al., 2004; Mukherjee et al., 2003; Prinz and Koenecke, 2012; Tang et al., 2012). The anti-Her2 ON NASCAR was introduced into Treg cells to evaluate whether GZV can regulate Treg activity. We incubated these cells with anti-CD19 CAR-expressing CD4+ T cells and Her2+/CD19+ NALM6 cells in the absence and presence of GZV (**Figure S3A**). CD4+ proliferation was evaluated by staining with a fluorescent cell tracer and tracking proliferation following 6 days of incubation. Resulting data showed that ON NASCAR Tregs were able to activate (via CD69 levels) in the presence of GZV (**Figure S3B**) and suppress CD4+ CAR T cell activation at a level comparable to that of Tregs expressing a traditional anti-Her2 CAR (**Figure S3C and S3D**).

### OFF NASCAR

To complement the ON NASCAR system, we also devised strategies to inhibit CAR signaling activity in response to NS3 inhibition. An OFF NASCAR was constructed using a 2-component split-polypeptide configuration similar to previously described drug-controllable receptor systems (Wu et al., 2015). The first component contains the scFv, a transmembrane domain, and an NS3-binding peptide. A second, membrane-tethered component contains the DAP10 ectodomain, NS3 domain, and CD3ζ signaling domain **(Figure 2A and Figure 4SA)**. By incorporating a catalytically-dead version of the NS3 domain (S139A), we are effectively using NS3 as an affinity domain. In the absence of GZV, the NS3-targeting peptide is expected to bind to NS3, bringing the two components together to drive CAR signaling. Upon GZV addition, the inhibitor competes with the peptide for binding to the NS3 substrate recognition site, thus facilitating dissolution of the CAR complex (**Figure 2A**). As before, we tested various designs to identify optimal configurations. Specifically, we designed four permutations (first component: a-d, second component: i-iv) of each component (**Figure S4A**). In these variations, the NS3-binding peptide sequence was positioned C-terminally either to the hinge region or to the individual costimulatory domains in the first component, and the NS3 placed at various positions in the second component.

**Figure 2:**
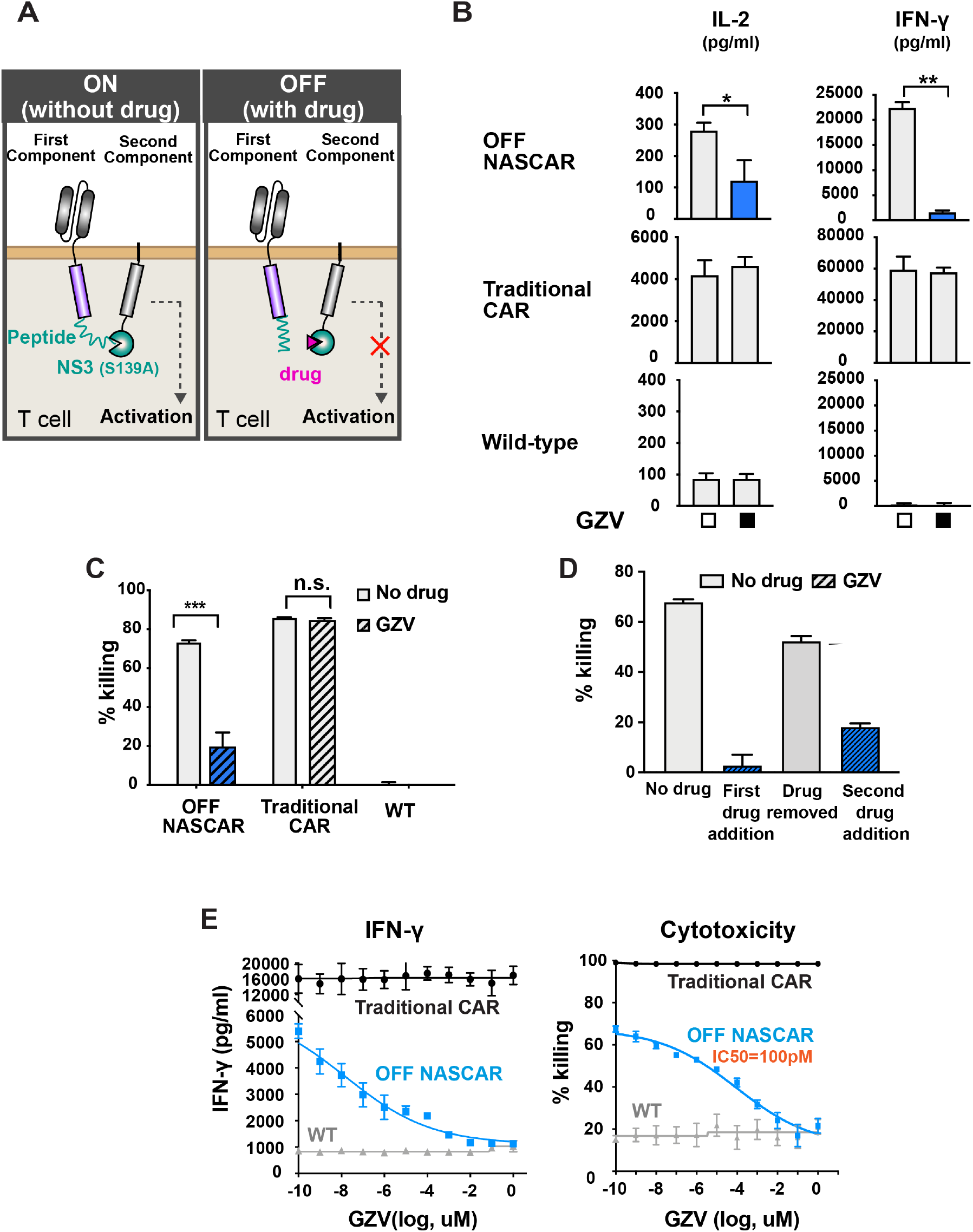
Characterization of the OFF NASCAR. **A**) Schematic of the OFF NASCAR mechanism. **B**) Primary T cell lines were treated with or without GZV (all in the presence of target cells), and cytokine levels quantified (mean ± s.d., n = 3, \**P* < 0.05, \*\**P* < 0.01 and \*\*\**P* < 0.001). **C**) Comparison of cytotoxicity levels of the OFF NASCAR with traditional CAR and wild-type cells (mean ± s.d., n = 3, \**P* < 0.05, \*\**P* < 0.01 and \*\*\**P* < 0.001). **D**) Cytotoxicity response of OFF NASCAR T cells to GZV being added, washed out and re-introduced after two days (mean ± s.d., n = 3). **E**) GZV dose response profile of various cell, as measured in cytokine levels and cell killing (mean ± s.d., n = 3).

We tested the performance of each configuration in Jurkat T cells first, with CD19 again used as the target of these CARs (**Figure S4C**). These Jurkat T cells had been modified to express GFP upon activation of the NFAT pathway, thus allowing us to detect T cell activation through GFP expression (Lin and Weiss, 2003). CAR-expressing Jurkat T cells were incubated with CD19+ NALM6 cells in the presence or absence of GZV and the resulting GFP levels measured. We found that some of the first component permutations (components a and b) had weak expression and therefore did not induce the NFAT transcription reporter (data not shown). However, first component permutations that contain CD28 (components c and d) resulted in CARs that could be activated when stimulated with target cells, and shut off when GZV was present. NFAT activity and CD69 expression were generally higher if both CD28 and 4-1BB costimulatory domains were in the first component (attached to the scFv), and the combination with a second component consisting of 4-1BB followed by NS3 and CD3ζ (component i) allowed for even stronger T cell activation (**Figure S4C**). When paired with components c (scFv-CD28-peptide) and d (scFV-CD28-41BB-peptide), second components i (DAP10-41BB-NS3-CD3ζ) and iii (DAP10-NS3-CD3ζ) resulted in the highest CD69 fold change, and thus we proceeded with component combinations c+i, c+iii, d+i and d+iii in further analysis with human primary T cells (**Figure S4D**).

Among the variations tested in primary T cells, component c resulted in higher T cell activation (as measured in cytokine levels) in the absence of GZV than component d. Of the two variations incorporating component c, version c+i of the OFF CAR was able to shut off T cell activity most effectively. This version of the OFF NASCAR was thus chosen as our final design (**Figure S4B**). The OFF NASCAR has similar basal activity as unmodified T cells and comparable induced activity to traditional CAR-expressing T cells (**Figure 2B, 2C**). Moreover, varying the concentration of GZV indicated that we could regulate the level of T cell activity in a dose-dependent manner, with an IC50 of around 100 pM **(Figure 2E),** which is >200-fold lower than the ON NASCAR. The reusability of the OFF NASCAR was additionally tested in a cell killing assay by adding, removing, and re-adding GZV, with results indicating that the OFF NASCAR is able to respond a second time following previous stimulation **(Figure 2D)**.

### NASCARs in xenograft leukemia model

After confirming ON and OFF NASCAR activity *in vitro*, we proceeded to test their functionality using anti-CD19 NASCARs in a blood tumor model commonly used as a preclinical model for evaluating CD19 CARs(Barrett et al., 2011). For mice receiving the NASCARs, GZV was dosed every day at 25 mg/kg for 14 days. Tumor growth was monitored via IVIS imaging of luciferase from NALM6 cancer cells over the course of 28 days (**Figure 3A**). As a control, it was verified that a daily injection of GZV of up to 25mg/kg for 14 days was not toxic to the NSG mice, nor did it reduce tumor growth (**Figures S5A-D**).

**Figure 3:**
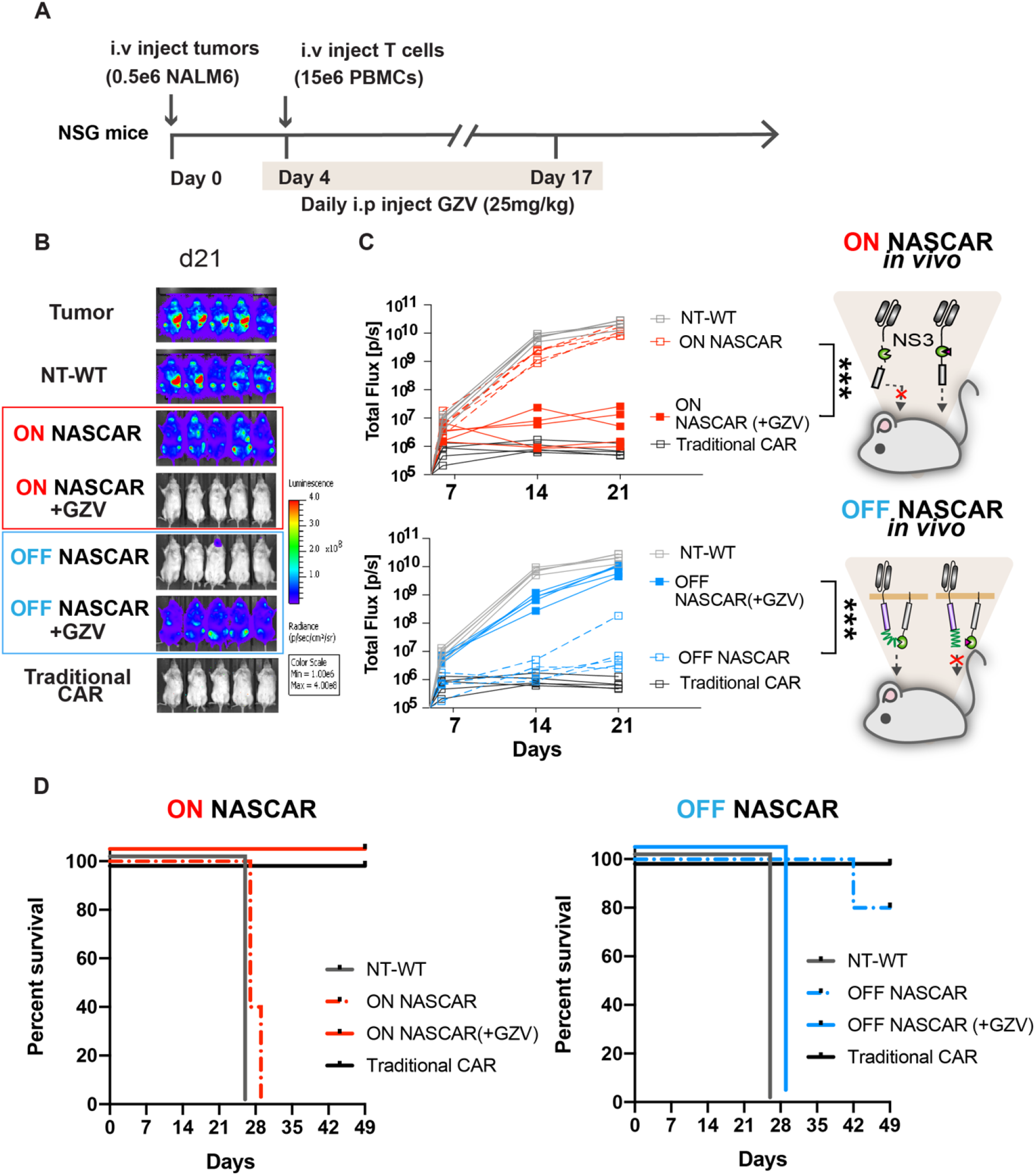
ON and OFF NASCAR are functional in a mouse xenograft tumor model. A) Timeline of *in vivo* experiments. B) IVIS imaging of groups treated with (1) no T cells, (2) non-transduced T cell (NT-WT), (3) ON NASCAR T cells, (4) ON NASCAR T cells with GZV, (5) OFF NASCAR T cells, (6) OFF NASCAR T cells with GZV, or (7) Traditional CAR T cells by day 21. C) Tumor burden was quantified as total flux (photons/s) from the luciferase activity of each mouse using IVIS imaging (n = 5, \**P* < 0.05, \*\**P* < 0.01 and \*\*\**P* < 0.001). D) Kaplan-Meier survival curves for the various mice treatment groups.

The appropriate working concentration of GZV was determined by testing the following concentrations of GZV and dosing durations on mice receiving the ON NASCAR: 50 mg/kg for 7 days, 25 mg/kg for 14 days, and 10 mg/kg for 14 days. It was found that 25 mg/kg of GZV for a duration of 14 days was sufficient for controlling ON NASCAR activity and eradication of tumor cells. Conversely, tumor regrowth was observed in mice receiving 50 mg/kg GZV for 7 days, possibly due to the shorter duration of drug administration (**Figure S5E**). As *in vitro* data indicated that the IC50 of GZV for the OFF NASCAR is lower than that of the ON NASCAR, it was presumed that a dose of 25 mg/kg GZV should be sufficient for controlling OFF NASCAR activity as well.

The ON and OFF NASCARs were then tested *in vivo* for their responsiveness to GZV and ability to clear tumors. We observed that mice injected with ON NASCAR T cells and treated with GZV were able to fully clear the tumor within 28 days, while those that did not receive GZV bore high tumor burdens. Additionally, mice receiving OFF NASCAR T cells cleared tumors in the absence of GZV, but displayed increased tumor burden when treated with GZV (**Figures 3B-C and Figures S6A-B**). We observed 0% survival in the group of mice that had not received GZV with the ON NASCAR, and those that had received GZV with the OFF NASCAR. In contrast, of the mice receiving the ON NASCAR and GZV, or the OFF NASCAR without GZV, 100% and 80% of them survived until day 49 respectively (**Figure 3D**). Overall, these results demonstrate that the ON and OFF NASCARs are functional in an *in vivo* leukemia model.

### Dual-gated NASCAR circuits

To explore more advanced inducible CAR designs, the ON NASCAR was first used to create an AND gate. A dual-gated and tunable logic AND gate could enhance the targeting specificity of CAR-T cells. To accomplish this, we used an orthogonal inducible technology involving the dihydrofolate reductase (DHFR) degron (DD). DHFR reduces dihydrofolate to tetrahydrofolate and is essential for *E. coli* replication and survival, and trimethoprim (TMP) is an antibiotic that can bind to *E. coli* DHFR with a much higher affinity than its mammalian counterpart. Fusion of the engineered DHFR domain with a protein of interest destabilizes the protein(Iwamoto et al., 2010). TMP binding to the DHFR domain stabilizes the fusion protein, thus serving as an inducible protein expression system. Here, we generated a combinatorial logic AND gate by distributing CAR intracellular signaling domains between two distinct scFvs, with each signaling domain associated with either the NS3 or DD. Our AND gate NASCAR system comprised of an anti-Her2 receptor with the NS3 domain positioned between the scFv and CD3ζ domains, followed by a second anti-Axl receptor with the DD positioned after the CD28 domain, thus generating a functional dual-molecule controllable CAR (**Figure 4A**).

**Figure 4:**
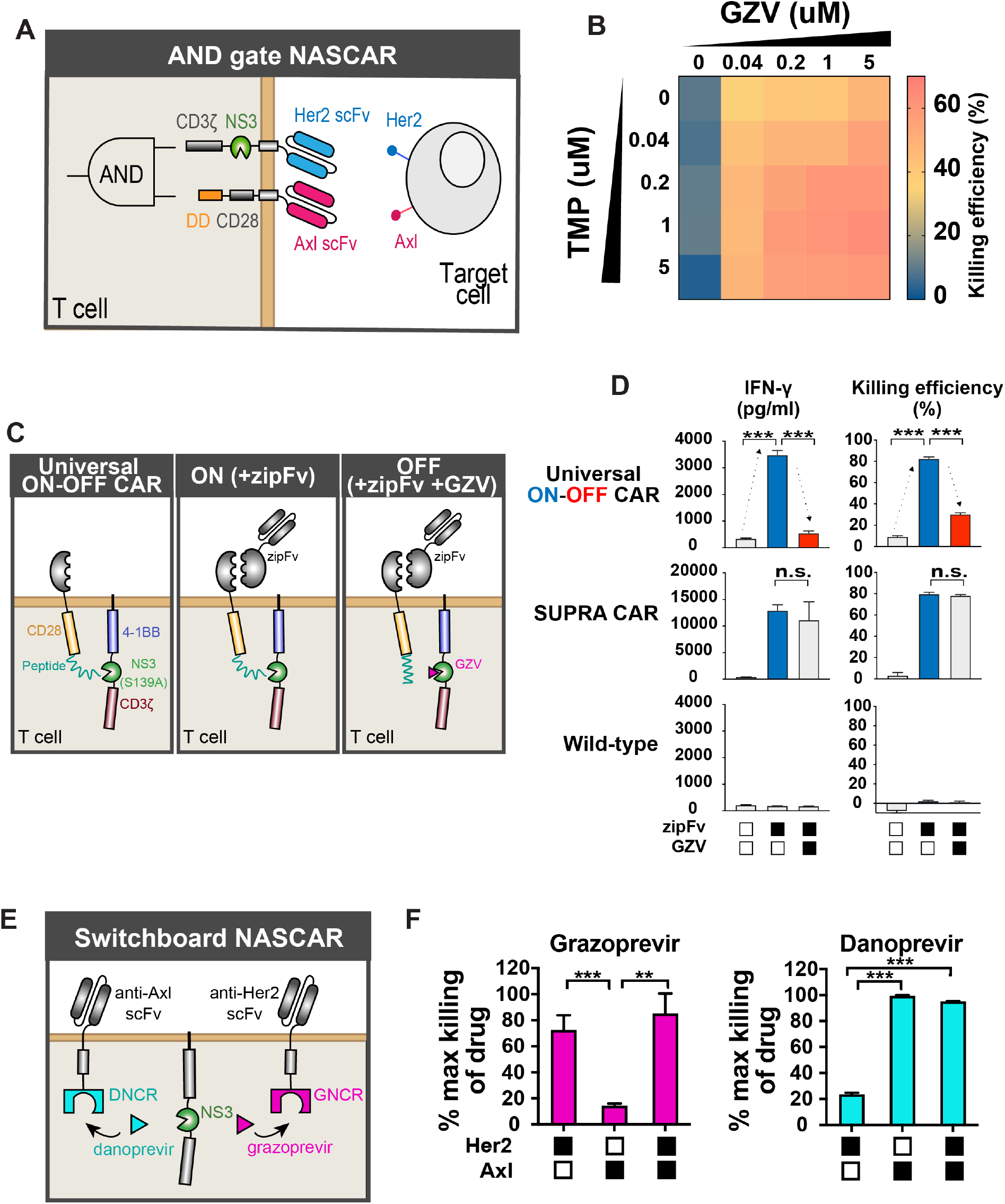
NS3 domains enable the design of dual gated NASCAR circuits. **A**) Schematic of the AND gate NASCAR that incorporates an ON NASCAR and a distinct DD CAR. **B**) Dose response of the AND gate NASCAR T cell killing in response to increased levels of GZV and TMP (n = 3, mean values are displayed). **C**) Schematic the universal ON-OFF NASCAR mechanism. **D**) Comparison of universal ON-OFF NASCAR T cell activity with traditional CAR and wild-type T cells, treated with combinations of zipFv and GZV, as measured by IFN-γ levels and cell killing (mean ± s.d., n = 3, \**P* < 0.05, \*\**P* < 0.01 and \*\*\**P* < 0.001). **E**) Schematic of switchboard NASCAR mechanism. **F**) Cytotoxicity levels of switchboard NASCAR in primary T cells when treated with various antigen-expressing target cells, normalized to the maximum killing observed under each drug (mean ± s.d., n = 3, \**P* < 0.05, \*\**P* < 0.01 and \*\*\**P* < 0.001).

The AND gate NASCAR was expressed in CD4+ T cells and CD8+ T cells separately (**Figure S7A**), and the drug tunability of the system tested. To activate the AND gate NASCAR, we co-cultured these T cells with Her2+/Axl+ NALM6 cells and varied the amount of GZV and TMP. Results indicated that cytotoxicity could be modulated by varying the amounts of the small molecules in CD8 T cells, although basal killing was observed in the presence of solely GZV (**Figure 4B and Figure S7B**). This could stem from low activation of the NS3-containing component, which forms a first-generation CAR. First generation CARs activate T cells to a weaker extent without the costimulatory domains, as compared to second or third generation CARs, but could still be sufficient for triggering this basal level of cytotoxicity *in vitro* (Brocker and Karjalainen, 1995; Hombach and Abken, 2013; Imai et al., 2004). Modulation of CD69 and cytokine levels was also observed by varying the amounts of the small molecules, suggesting that the level of T cell activation could be tuned using different doses of the two drugs in CD4+ T cells (**Figure S7C**).

In addition to the AND gate NASCAR, we also designed controllable CARs that can change antigen targets. These drug-gated CARs are useful for combating relapse caused by antigen escape. We created a universal OFF CAR by applying the OFF NASCAR design to existing SUPRA CAR technology. The SUPRA CAR consists of two components: a universal zipCAR that is formed by the fusion of a leucine zipper and intracellular CAR signaling domains, and an adaptor protein called zipFv that is formed by the fusion of an scFv and the cognate leucine zipper. When the zipFv bridges the zipCAR T cells and target cancer cells, it allows for activation through the zipCAR. To create our universal ON-OFF NASCAR, we divided the intracellular CAR domains into two components, similar to the design of the OFF NASCAR, and replaced the scFv with a leucine zipper system similar to that of the SUPRA CAR. In the absence of zipFv or GZV, the CAR remains inactive as there is no scFv targeting it to any cell. However, when zipFv is added, the zipCAR can be activated by target cells. This activation is then inhibited when GZV outcompetes the NS3-specific peptide for binding to the catalytically-dead NS3 domain (**Figure 4C**). This universal ON-OFF NASCAR was introduced into primary CD8+ T cells, and through IFN-γ production measurements we found that these cells were able to switch ON in the presence of anti-Her2 zipFv and target Her2+ NALM cells, and shut OFF when GZV was present. Cytotoxicity of these T cells was additionally measured in a killing assay involving Her2+ NALM6 target cells, and results recapitulated what was observed with the cytokine data (**Figure 4D**). Furthermore, we found that the level of activation obtained by cells expressing this ON-OFF NASCAR could be adjusted by varying zipFv concentration, demonstrating a high level of tunablity (**Figure S8)**.

Lastly, NS3 “reader” proteins were applied to CAR design to generate a switchboard NASCAR circuit, which allows it to switch to a different target antigen through GZV or DNV administration. These “reader” proteins bind specific inhibitor-bound states of NS3, and are expected to behave orthogonally(Foight et al., 2019). To develop the switchboard NASCAR circuit, we adopted the DNV/NS3 complex reader (DNCR) and GZV/NS3 complex reader (GNCR), which are regulated by DNV and GZV respectively. Similar to the OFF NASCAR, the NS3 reader CARs were designed by splitting the CAR into two components. The first component consists of the scFv, CD28 costimulatory domain, and reader domain, while the second component encodes a catalytically-dead NS3 and the CD3ζ signaling domain. The NS3 domain binds to the GNCR in the presence of GZV, and the DNCR in the presence of DNV, allowing different scFvs to bind to the CD3ζ depending on the drug added (**Figure 4E**). We first tested the function of our DNCR and GNCR CARs separately in cells using an anti-Axl and anti-Her2 CAR respectively. Primary T cells were transduced with lentivirus encoding the reader CARs and incubated with or without inhibitors, and NALM6 target cells expressing Her2, Axl, or both antigens. Both reader CARs showed an increase in cytotoxicity and cytokine release levels when their respective inducer was present, along with the target cell line expressing their specific antigen (**Figures S8A and S8B**). With both reader CARs functional, we proceeded to express both CARs within the same cell. While the GNCR CAR indicated lower activity than the DNCR CAR, these dual CAR-expressing cells indicated increased cytokine levels and killing of Her2+ target cells when GZV was present, and increased cytokine levels and killing of Axl+ target cells when DNV was present (**Figures 4F and S8C**). This suggests that we are able to switch CAR antigen specificity using these two drugs.

## DISCUSSION

### Clinically-approved drug gated CAR circuits

The need to address the safety of CAR-T cell therapy is underscored by its potentially fatal side effects(Rafiq et al., 2020). Indeed, toxicity from CD19 directed CAR-T cells has been observed and best characterized in clinical trials. Although CAR T cells represent a revolutionary approach to cancer treatment, strategies to circumvent their potential lethality are needed. To address this, increased control over CAR T cell activity is required, and drug-inducible strategies to regulate the behavior of engineered cellular agents following their infusion into the body are especially desirable. While several drug-inducible CAR technologies currently exist that do so, none yet are reliant on a clinically-approved pharmaceutical agent with a favorable safety profile. Here, we have exploited the NS3 cis-protease domain and its corresponding inhibitors to devise a chemogenetic toolset for regulating the timing and magnitude of CAR T cell activities. Features that distinguish these systems from previously reported designs are their ability to respond to clinically-approved anti-viral compounds, and versatility with which they can be used to generate on and off signaling systems. In the work described above, we demonstrated an ON switch, OFF switch, and three complex CAR circuits (AND-gate, universal ON-OFF, and switchboard NASCARs) in which NASCARs were combined with other well-characterized inducer responding domains in order to achieve regulable and safe CARs. Each of these CARs has a unique design and can be regulated effectively by clinically-approved drugs. We have further demonstrated the efficacy of the ON and OFF NASCARs *in vivo* using a xenograft mouse model. The use of FDA-approved drugs as inducers in these technologies is beneficial as it can accelerate their clinical use. Furthermore, the tunability of the ON and OFF NASCARs using these drugs allows for the tailoring of T cell activation levels to individual patients’ needs. We envision these NASCARs T cell immunotherapy will be able to, in real time, modulate T cell activity to mitigate complications like CRS and off-target activities. Additionally, as exhibited with the ON CAR, we have shown the potential of these systems to target different antigens (anti-CD19 and anti-Her2) and operate in different cell types (CD4+, CD8+, and regulatory T cell), which suggests the applicability of these NASCARs to a broad range of cancer and cell types.

### Engineering advanced inducible CAR circuits to meet the current demands of immunotherapy

In addition to ON and OFF switches, we have demonstrated how the combination of different technologies can create more complex CAR circuits with expanded and necessary capabilities. Among the current drawbacks of T cell immunotherapy are off-target activities and antigen escape. One of the ways to solve these issues is to increase the specificity for tumor antigens through multiple-antigen sensing or changing the target antigen when desired. Combinatorial antigen sensing through AND-gate logic allows T cells to activate only in the presence of multiple antigens, which addresses problems that may arise when low levels of a targeted antigen are expressed on healthy tissue. We have developed our dual-gated AND gate CAR by combining NS3 technology with an existing degradation domain. This AND-gate logic is controlled through the use of two orthogonal drugs (GZV and TMP), which adds an additional layer of safety on top of the multiple-antigen sensing. We have further developed a universal ON-OFF NASCAR. To do this, we configured the SUPRA CAR, which can target different antigens by switching the target zipFv, with our OFF NASCAR technology to immediately inhibit CAR function when desired. The universal ON-OFF NASCAR is not only able to mitigate antigen escape by changing with zipFv of interest when desired, but also incorporates a safety switch that shuts off activity immediately when needed. Lastly, the compatibility of several FDA-approved drugs with the NS3 domain is used to our advantage in the design of our switchboard CAR. Using proteins that recognize specific inhibitor-bound states of the NS3, the switchboard CAR is able to change target antigens by simply exposing the cells to different drugs. It provides an additional method for attacking target cancer cells in the event of antigen escape. Applying the associated technologies of the NS3 domain to other synthetic biology tools allows us to further the possibilities of CAR T cell therapy. As the field of T cell immunotherapy advances, we foresee these capabilities as becoming increasingly important.

### The promise of viral proteases as synthetic biology tools in engineering T cells

The NS3 domain has been largely studied as a target for HCV treatment. Yet, we have expanded on its potential for use in synthetic biology applications, specifically in the design of CARs. Its uses as a self-cleaving entity and CID make it advantageous in receptor design due to its versatility. A further advantage of using viral proteases in CAR design is the well-studied, validated, and robust collection of clinically-approved viral inhibitors in the form of small molecules or peptides that have already been developed. Other viral proteases, such as the SAR-CoV2 MPro(Jin et al., 2020; Sacco et al., 2020), have been targeted as key entities for developing inhibitors to treat the viral infection. The NS3 protease and its inhibitors utilized here are a prime example of how they can be integrated into CAR design, providing clinically-ready tools for enhancing the safety and efficacy of CAR-T cell therapy. Given the availability of multiple NS3-targeting drugs, and due to the continued development of new inhibitory compounds, this research provides a streamlined framework for the translation, elaboration, and adaptation of antiviral strategies into effective therapeutic platforms for the treatment of life-threatening, non-viral malignancies.

## ACKNOWLEDGEMENTS

W.W.W. acknowledges funding from the Boston University Ignition Award and NIH Director’s New Innovator Award (1DP2CA186574). A.S.K. acknowledges funding from the NIH (R01EB029483), NIH Director’s New Innovator Award (1DP2AI131083), and DARPA Young Faculty Award (D16AP00142). J.T.N and E.P.T were supported by NIH NIGMS grant (R35-GM128859-01), as well as by the Reidy Family Career Development Award. E.P.T. was supported through a T32 NIH Training Grant awarded to Boston University (EB006359). We also thank Wong lab members for suggestions on the manuscript; Dr.Todd Blute from the BU Proteomics & Imaging Core Facility for flow cytometry assistance; BU IVIS imaging core facility for mice *in vivo* imaging.

## AUTHOR CONTRIBUTIONS

H.S.L and N.M.W designed and generated genetic constructs, performed experiments, analyzed the data, and generated figures. E.T. and J.T.N. helped design NASCARs. A.S.K supervised the project and analyzed the data. W.W.W conceived and supervised the project and analyzed the data. All authors commented on and approved the paper.

## MATERIALS AND METHODS

### KEY MATERIALS

| REAGENT or RESOURCE | SOURCE | IDENTIFIER |
| --- | --- | --- |
| **Antibodies** |  |  |
| APC-Cy7-anti-human CD69 | BD Pharmingen | 557756 |
| FITC Mouse Anti-Human CD4 Clone M-T477 | BD Biosciences | 556615 |
| Pacific Blue™ Mouse Anti-Human CD8 | BD Biosciences | 558207 |
| Dynabeads™ Human T-Activator CD3/CD28 | Thermo Fisher Scientific | 11132D |
| Alexa Fluor 647 anti-V5 Tag | Invitrogen | 451098 |
| Alexa Fluor 488- anti-myc Tag | R&D Systems | IC3696G |
| **Cell Culture Reagents** |  |  |
| RPMI-1640 | Lonza | 12-702Q |
| Ultraculture | Lonza | 12-725F |
| DMEM | Corning | 10-013 |
| X-VIVO 15 | Lonza | 04-418Q |
| Human AB serum | Valley Medical | HP1022 |
| FBS | Thermo Fisher Scientific | 10437-028 |
| 2-Mercaptoethanol | Thermo Fisher Scientific | 31350010 |
| L-glutamine | Corning | 25-005-CL |
| Penicillin/Streptomycin | Caisson | PSL01 |
| Sodium Butyrate | Alfa Aesar | A11079 |
| **Chemicals, Peptides, and Recombinant Proteins** |  |  |
| Grazoprovir | MedChemExpress | HY15298 |
| Grazoprovir potassium salt | MedChemExpress | HY-15298A |
| Danoprevir | ApexBio | A4024 |
| Glecaprevir | MedChemExpress | HY-17634 |
| Simeprevir | MedChemExpress | HY-10241 |
| Boceprevir | MedChemExpress | HY-10237 |
| TMP | Sigma-Aldrich | 92131 |
| XenoLight D-Luciferin - K+ Salt Bioluminescent Substrate | Perkin Elmer | 122799 |
| N-Acetyl-L-cysteine | Sigma-Aldrich | A9165 |
| Lenti-X™ Concentrator | Clontech | 631232 |
| RetroNectin® Recombinant Human Fibronectin Fragment | Clontech | T100B |
| Phosphate-Buffered Saline (10X) pH 7.4 | Thermo Fisher Scientific | AM9625 |
| PEG8000 | Thermo Fisher Scientific | BP233-1 |
| Sodium Chloride | Thermo Fisher Scientific | AAJ2161836 |
| DMSO | CHEM-IMPEX INT'L INC | 00635 |
| PEG400 | MedChemExpress | HY-Y0873A |
| **Critical Commercial Assays** |  |  |
| RosetteSep™ Human CD4+ T Cell Enrichment Cocktail | STEMCELL Technologies | 15062 |
| RosetteSep™ Human CD8+ T Cell Enrichment Cocktail | STEMCELL Technologies | 15063 |
| EasySep™ Human CD4+CD127lowCD25+ Regulatory T Cell Isolation Kit | STEMCELL Technologies | 18063 |
| Human IL-2 ELISA Set | BD Biosciences | 555190 |
| Human IFN-γ ELISA Set | BD Biosciences | 555142 |
| XenoLight D-Luciferin - K+ Salt Bioluminescent Substrate | PerkinElmer | 122799 |
| CellTrace™ Violet Cell Proliferation Kit, for flow cytometry | Thermo Fisher Scientific | C34557 |
| RosetteSep™ Human CD4+ T Cell Enrichment Cocktail | STEMCELL Technologies | 15062 |
| **Experimental Models: Cell Lines** |  |  |
| HEK293FT | N/A | N/A |
| NFAT-Jurkat | Art Weiss (UCSF) | N/A |
| NALM6 (Luc+ BFP+) | This paper | N/A |
| NALM6 (HER2+ Axl+ Mesothelin+ Luc+) | This paper | N/A |
| NALM6 (HER2+ mCherry+) | This paper | N/A |
| NALM6 (Axl+ GFP+) | This paper | N/A |
| **Experimental Models: Organisms/Strains** |  |  |
| NOD.Cg-*Prkdc<sup>scid</sup> Il2rg<sup>tm1Wjl</sup>*/SzJ (female 4-6 weeks) | Jackson Laboratory | 005557 |
| **Recombinant DNA** |  |  |
| pHR-SFFV vector | Addgene | 79121 |
| PB-pCAG-MCS-EF1a-zeocine-BFP vector | This paper | N/A |
| **Software and Algorithms** |  |  |
| Graph Pad Prism 7 | Graph Pad | N/A |
| FlowJo V10 | TreeStar | N/A |
| Living Image | Perkin Elmer | N/A |

### CONTACT FOR REAGENT AND RESOURCE SHARING

Reagent requests should be directed and will be fulfilled by lead author Wilson Wong.

### EXPERIMENTAL MODEL AND SUBJECT DETAILS

#### Source of Primary Human T Cells

Blood was obtained from the Blood Donor Center at Boston Children’s Hospital (Boston, MA) as approved by the University Institutional Review Board.

#### Animal Model Details

Animal studies were conducted at the Boston University Medical School Animal Science Center under a protocol approved by the Boston University Institutional Animal Care and Use Committee. Female NSG mice, 6-8 weeks of age, were purchased from Jackson Laboratories (#005557) and were used for all *in vivo* mouse experiments.

#### CAR Constructs Design

All CAR constructs were introduced into primary human T cells and Jurkat T cells using pHR lentivectors, and plasmids were packaged into lentiviruses using pDelta, Vsvg and pAdv packaging and envelope plasmids(Zufferey et al., 1998). Expression of the CARs was driven by an EF-1 alpha promoter and kozak sequence. CARs contained either a myc or V5 tag after the scFv, or a mCherry fluorescent protein at the C-terminus in order to verify construct expression. In OFF NASCAR, Universal ON-OFF NASCAR experiments, both components were introduced into cells using a single construct, with the components separated by either a P2A or T2A sequence. The two components of the AND gate NASCAR and switchable NASCAR were introduced on separate constructs.

#### Cell culture

Lentiviruses were generated using HEK293FT cells, which were cultured in Dulbecco’s modified Eagle’s medium (DMEM) supplemented with 10% fetal bovine serum (FBS; Thermo Fisher, 10437028), penicillin/streptomycin (Corning, 30001CI), L-glutamine (Corning, 25005CI) and 1mM sodium pyruvate (Lonza, 13115E). Primary peripheral blood mononuclear cells (PBMCs), CD4+ and CD8+ T cells were isolated from whole peripheral blood obtained from healthy donors at the Blood Donor Center at Boston Children’s Hospital (Boston, MA) as approved by the University Institutional Review Board (IRB). CD4+ and CD8+ T cells were isolated using the RosetteSep Human T cell enrichment cocktail (STEMCELL Technologies, 15022 and 15023), and PBMCs were isolated using Lymphoprep (STEMCELL Technologies, 07851). Regulatory T cells were isolated using the EasySep human CD4+CD127lowCD25+ regulatory T cell isolation kit according to manufacturer’s protocol (STEMCELL Technologies, 18063). Primary T cells were cultured in X-Vivo 15 media (Lonza, 04-418Q) supplemented with 5% human AB serum (Valley Biomedical, HP1022), 10mM N-acetyl L-Cysteine (Sigma, A9165), 55μM 2-Mercaptoethanol (Thermo Fisher, 21985023), and 30-200U/ml IL-2 (NCI BRB Preclinical Repository). N-acetyl L-Cysteine and 2-mercaptoethanol were removed during the assay of regulatory T cell suppression experiment. Jurkat T cells and NALM6 cells were cultured in RPMI 1640 supplemented with 5% FBS, L-glutamine and penicillin/streptomycin. CD19-positive NALM6 cells expressing Her2, mCherry, and GFP, or just BFP alone, were used as target cells in assays.

#### Lentivirus generation and transduction

HEK293FT cells were co-transfected with lentivirus packaging/envelope plasmids (described above) and CAR-encoding vectors using polythylenimine (PEI). After 24 hours, media was replaced with ultraculture (Lonza, 12-725F) supplemented with penicillin/streptomycin, L-glutamine, 0.5M sodium butyrate and 1mM sodium pyruvate. Virus-containing media was collected for the following 48 hours, and spun down to remove cell debris. Viral media was then concentrated either using Lenti-X Concentrator (Takara, 631232) at a 3:1 ratio and incubated at 4°C overnight prior to centrifugation at 1500xg for 45 minutes at 4°C, or using Lentivirus concentration solution (40% PEG8000 and 1.5M NaCl) at a 3:1 ratio and incubated at 4°C overnight prior to centrifuga-tion at 1600xg for 60minutes at 4°C. To transduce primary T cells, lentiviral spinfection was performed. Cells were thawed 2 days prior to spinfection, and 1 day prior to spinfection cells were activated using Human T-activator CD3/CD28 Dynabeads (Thermo Fisher, 11131D) and cultured in 200U/ml IL-2. Concentrated virus was plated on non-TC treated 6-well plates coated with retronectin (Takara T100B), and spun for 90 minutes at 1200xg. Virus was then aspirated and activated T cells added to the virus-coated wells, followed by incubation at 37°C. To transduce Jurkat T cells, concentrated virus was diluted in RPMI 1640, mixed with cells and incubated for 72 hours at 37°C before virus was washed out.

#### Antibodies and cell dyes

To stain for CD69, an APC-Cy7-conjugated mouse anti-human CD69 antibody (BD Pharmingen, 557756) was used at a dilution of 1:100. Myc-tagged and V5-tagged CARs were stained using an Alexa Fluor 488-conjugated mouse anti-myc antibody (R&D Systems, IC3696G) and an Alexa Fluor 647-conjugated mouse anti-V5 antibody (Invitrogen, 451098) respectively. For surface staining, cells were washed twice with FACS buffer (1x phosphate buffered saline (PBS), 0.1% NaN3, 1% BSA, 2mM EDTA) before incubating with antibody in the dark at room temperature for 40 minutes. Cells were then washed and resuspended with FACS buffer before analysis on an Attune NxT flow cytometer.

#### Cytotoxicity assay

Cytotoxicity assays were carried out by incubating primary T cells and NALM6 target cells at a 1:1 effector to target (E:T) ratio overnight at 37°C. In the case of the switchboard NASCAR experiments, a 1:2 E:T ratio was used. 0.5-1μM GZV (MedChemExpress, HY-15298) was added at the time of assay, and DMSO was used as a control vehicle. When testing other NS3 inhibitors, various concentrations in the range of 0-5 μM of GZV, danoprevir, simeprevir, glecaprevir and boceprevir were used in dose response experiments. For dose response experiments for ON and OFF NASCAR, 0-1μM of GZV were served, and for AND gate NASCAR, 0-5μM of GZV and (or) 0-5μM of TMP were served. Following a 16hour incubation, the supernatant was saved and cells were analyzed by flow cytometry to count the number of remaining live NALM6 cells by gating for either GFP+/mCherry+ cells or BFP+ positive cells depending on the target NALM6 cell line used. Killing efficiency was calculated as the percentage of cells killed compared to control wells containing NALM6 cells and no T cells.

#### Cytokine release assay

Supernatant from cells incubated with various combinations of target cells and NS3 inhibitors was saved and tested in an enzyme-linked immunosorbent assay (ELISA) to measure IFN-gamma and IL-2 levels. The BD OptEIA human IFN-γ and IL-2 ELISA kits (BD Biosciences, 555142 and 555190) were used according to manufacturer’s instructions with a 0.05% Tween-20 in PBS (Thermo Scientific, 28352) wash buffer, and remaining reagents from BD OptEIA Reagent Set B (BD Biosciences, 550534). ELISAs were performed with 96-well MaxiSorp plates (Thermo Scientific, 442404).

#### Cell proliferation assay

Effector cells were stained with the Cell Trace Violet kit (Thermo Fisher, C34571) as per manufacturer’s instructions. Treg cells and Teff cells were mixed with NALM6 target cells expressing Her2 and Axl as target cells at Treg: Teff: Target cell = 2:1:1 ratio. 1μM GZV (MedChemExpress, HY-15298) was added at the time of assay and cells were collected for flow cytometry analysis after incubation for 5-6days.

#### Xenograft model

Female NOD.Cg-Prkdcscid Il2rgtm1Wjl/SzJ (NSG) mice, 6-8 weeks of age, were purchased from Jackson Laboratories (#005557) and maintained in the BUMC Animal Science Center (ASC). All protocols were approved by the Institutional Animal Care and Use Committee at BUMC.

In order to generate the intravenous blood tumor xenograft models, NSG (*NOD Cg-Prkdcscid Il2rgtm1Wjl/ SzJ)* mice were initially injected with 0.5 × 10^6^ luciferized Nalm-6 cells intravenously. After 4 days, CAR-T or NASCAR −T cells were infused intravenously. Grazoprevir potassium salts (MedChemExpress, HY-15298A) were dissolved in 2.5% DMSO, 30% PEG400 and 67.5% PBS and i.p. injected every day for indicated dosage and times. Tumor burden was measured by IVIS Spectrum (Xenogen) and was quantified as total flux (photons per sec) in the region of interest. Images were acquired within 10 minutes following intraperitoneal injection of 150mg/kg of D-luciferin (PerkinElmer #122799)

### QUANTIFICATION AND STATISTICAL ANALYSIS

Data between two groups was compared using an unpaired two-tailed t-test. All curve fitting was performed with Prism 7 (Graphpad) and p values are reported (not significant = p > 0.05, * = p < 0.05, ** = p < 0.01, *** =p < 0.001). All error bars are represented either SEM or SD.

---

**Figure S1:**
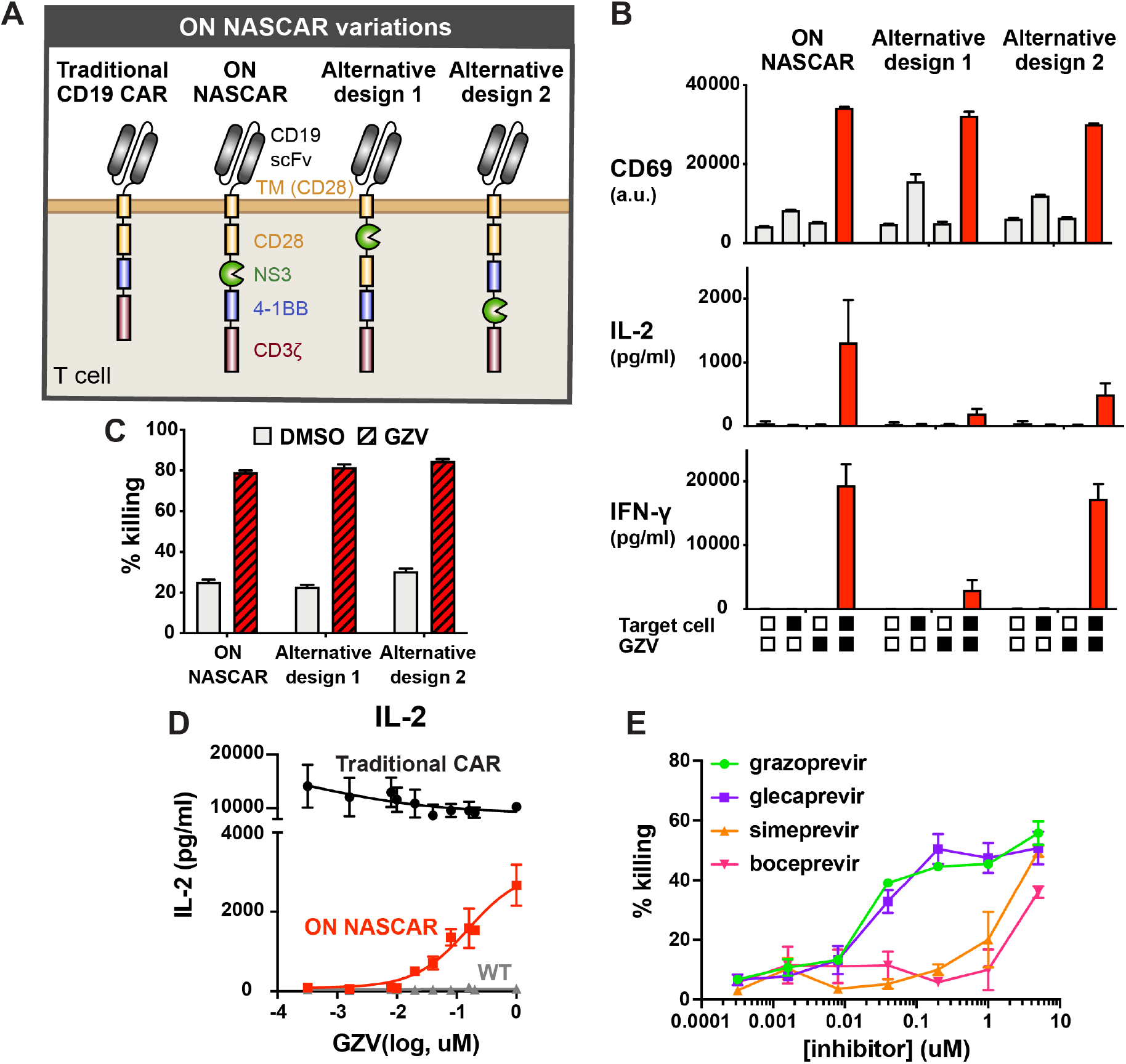
Variations of the ON NASCAR designs. **A**) Schematic of the variations of ON NASCAR designs. **B**) Regulation of the activity of ON NASCARs in primary T cells (measured by CD69 and cytokine levels) using combinations of target NALM6 cells and GZV. **C**) Cell killing of target NALM6 cells using primary T cells expressing ON NASCARs in the absence and presence of GZV. **D**) GZV dose response profile of NASCAR in primary T cell. **E**) Cytotoxicity of ON NASCAR-expressing primary T cells in response to different NS3 inhibitors.

**Figure S2:**
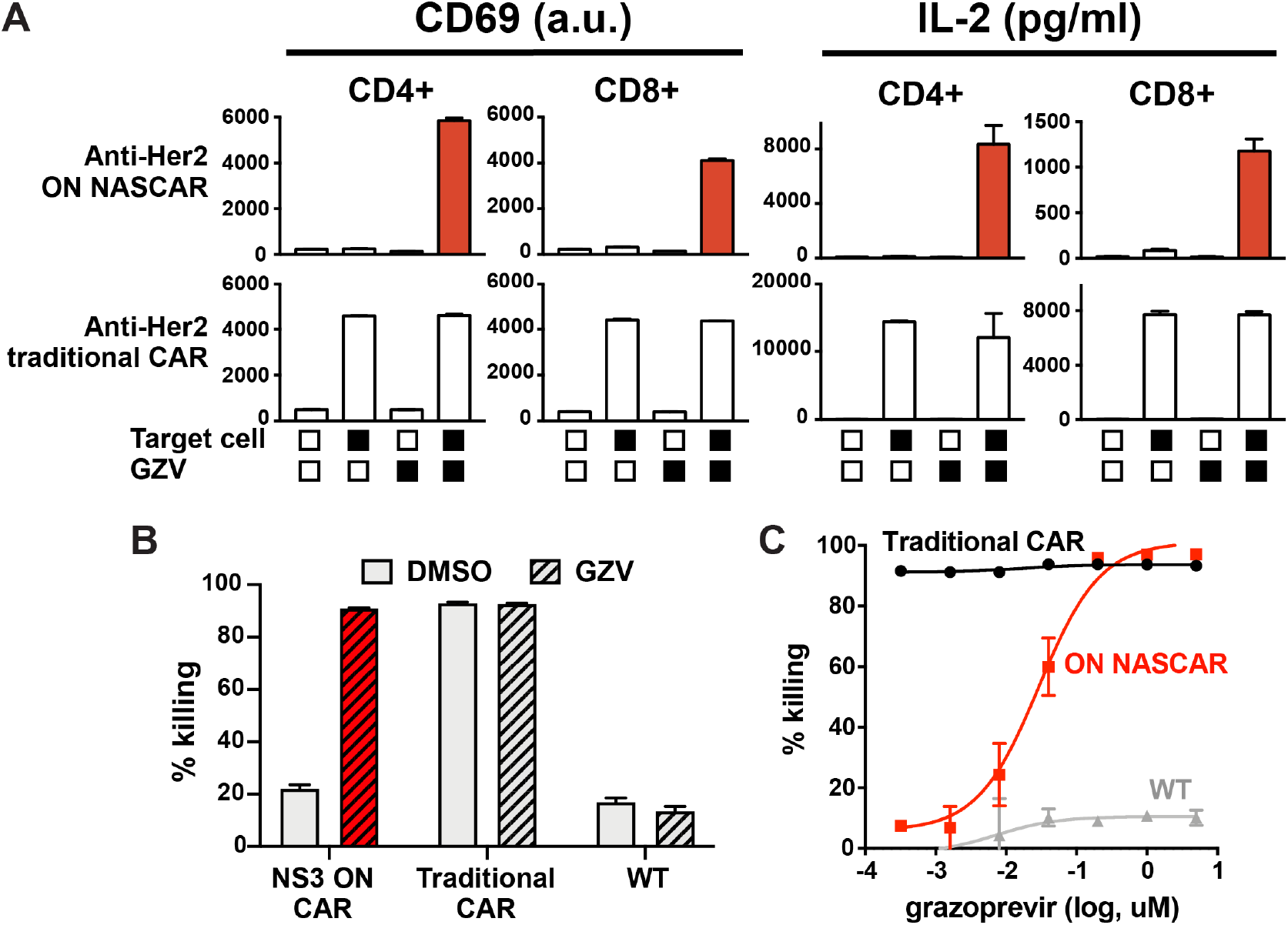
Application of ON NASCAR to a Her2 scFv. **A**) Response of the anti-Her2 ON NASCAR to combinations of target cells and GZV when expressed in CD4+ and COB+ T cell subsets. **B**) Cell killing ability of COB+ T cells expressing various anti-Her2 ON NASCAR, traditional anti-Her2 CAR, or no CAR. **C**) Dose response of anti-Her2 ON NASCAR COB+ T cells to increased amounts of GZV, as measured in cell killing efficiency.

**Figure S3:**
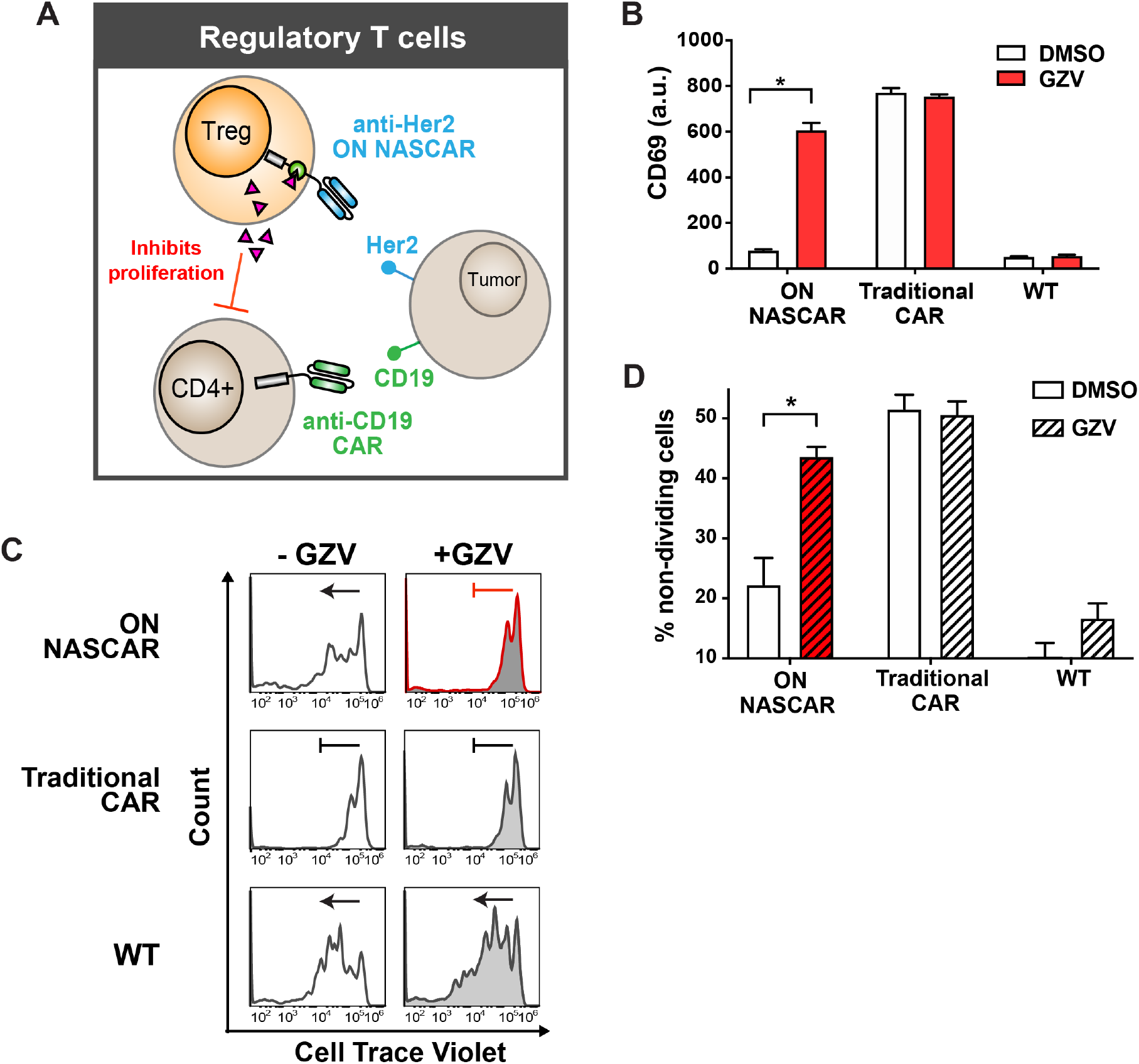
Application of ON NASCAR to regulatory T cells. **A**) Schematic of how ON NASCAR-expressing Treg cells interact with CD4+ T cells and target NALM6 cells in proliferation assay. **B)** Levels of early activation marker (CD69) expressed by ON NASCAR regulatory T cells in the presence and absence of NS3 inhibitor, compared with traditional CAR-expressing cells, and cells with no CAR. **C)** CD4+ T cell proliferation when co-incubated with various regulatory T cell lines in the presence and absence of GZV. **D)** Quan­ tifica-tion of non-dividing CD4+T cells from histogram proliferation data (Figure S3C).

**Figure S4:**
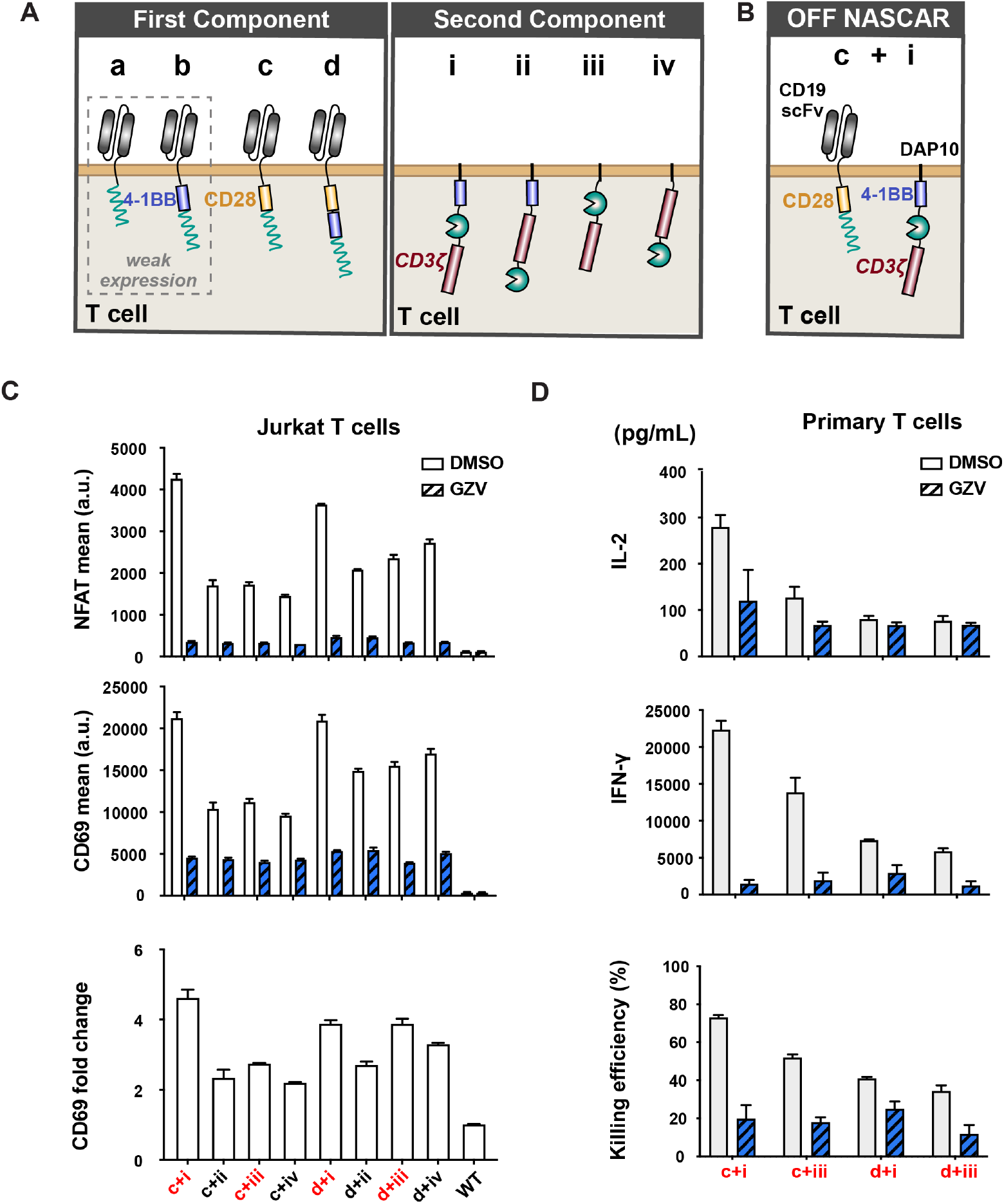
Variations on the OFF NASCAR designs. **A**) Variations on the two components that make up the OFF NASCAR. **B)** Schematic of the OFF NASCAR design. **C)** Jurkat T cell activity as measured by NFAT and CD69 levels when variations of the OFF NASCAR are expressed. Fold change observed in CD69 levels when GZV is added to Jurkat T cells expressing variations of the OFF CAR (mean ± s.d., n = 3). **D)** Cytokine release and cytotoxicity of primary T cells expressing different versions of the OFF NASCAR (mean ± s.d., n = 3).

**Figure S5:**
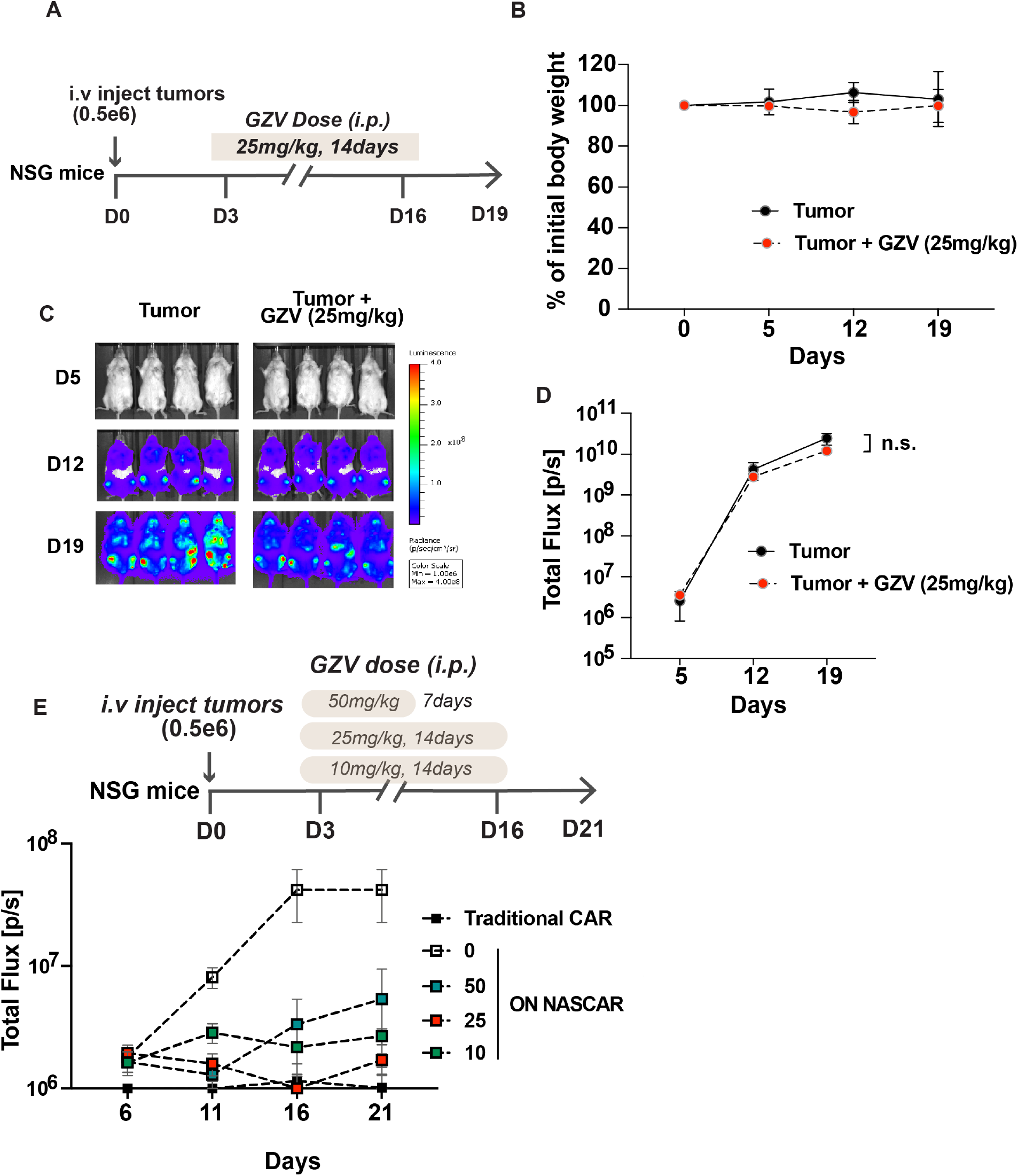
GZV dose optimization in vivo. **A**) Timeline for experiment testing the effect of GZV alone on tumor growth in vivo. **B)** Percentage of body weight to initial weight (day 0) was measured at days 5, 12 and 19 (mean ± s.d., n = 4). **C)** Luciferase levels from tumors imaged by IVIS for groups treated with (1) tumor alone and (2) tumor with GZV (25mg/kg for 2 weeks) at days 5, 12 and 19. **D)** Tumor burden was quantified as the total flux (photons/s) from the luciferase activity of each mouse using IVIS imaging (n = 4, mean ± sem). **E**) Top: Timeline for experiment testing variable GZV doses and administration duration for the ON NASCAR in vivo. Bottom: Tumor burden was quantified as total flux (photons/s) from the luciferase activity of each mouse using IVIS imaging (mean ± sem, n=4).

**Figure S6:**
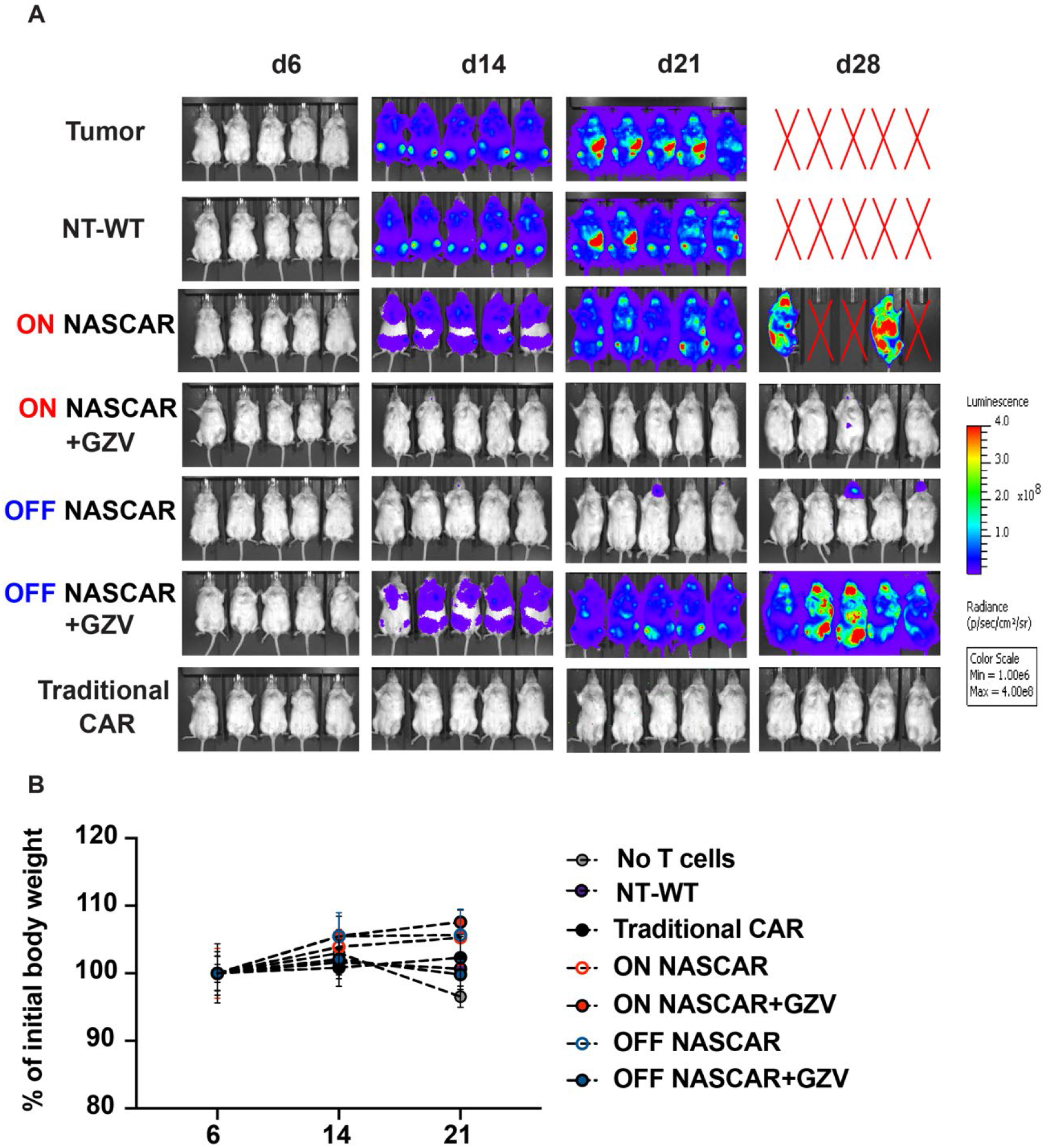
In vivo demonstration of ON and OFF NASCARs. **A)** IVIS imaging of groups treated with (1) no T cells, (2) non-transduced T cell (NT-WT), (3) ON NASCART cells, (4) ON NASCART cells with GZV, (5) OFF NASCAR T cells, (6) OFF NASCART cells with GZV, or (7) Traditional CART cells at days 6, 14, 21 and 28. **B)** Percent­ age of body weight to initial weight (day 0) was measured at days 6, 14 and 21. (mean ± s.d., n = 5)

**Figure S7:**
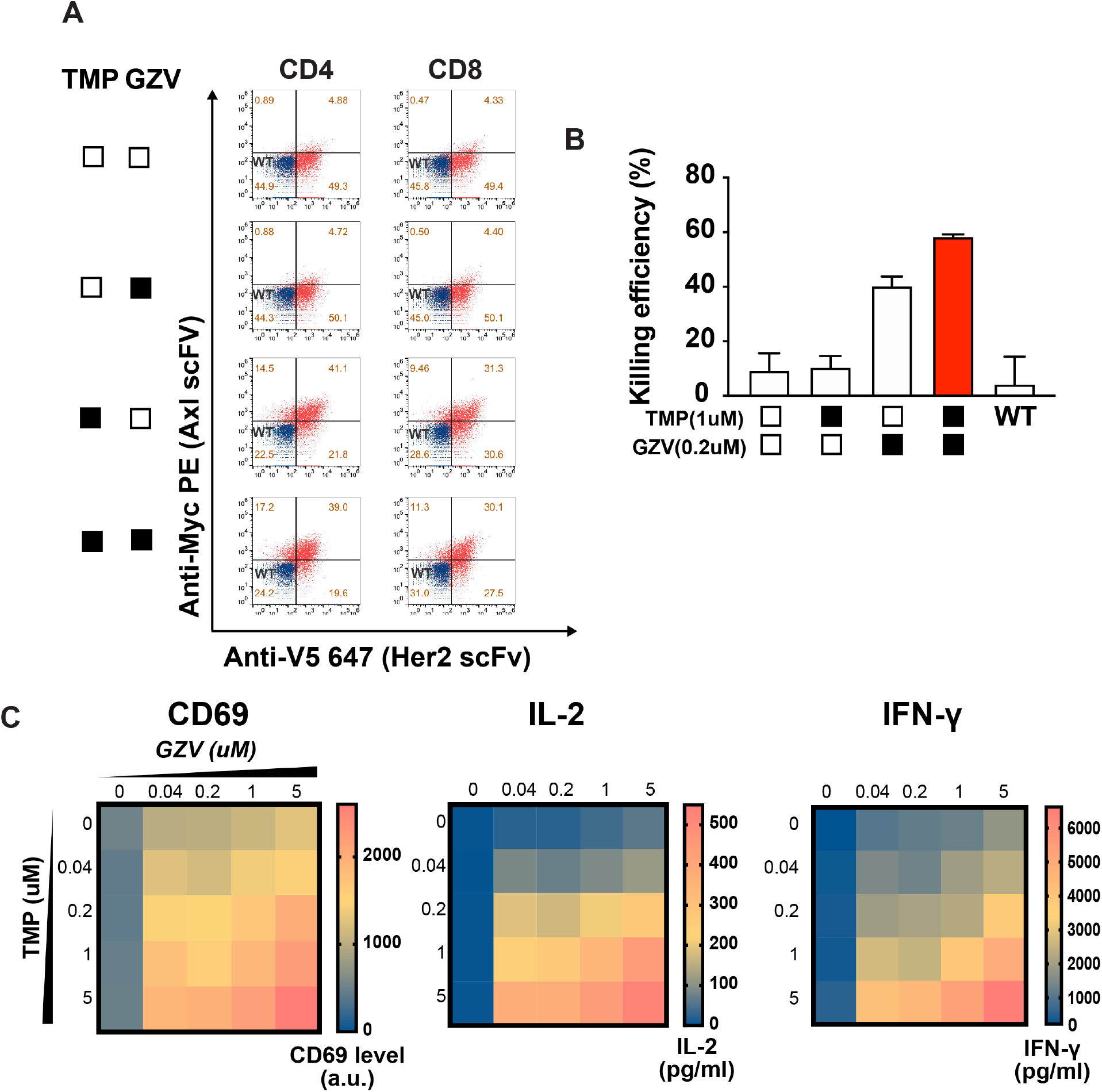
Cytotoxicity of cells expressing the AND gate NASCAR. **A)** Following treatment with combinations of TMP and GZV, anti-Her2 scFv and anti-Axl scFv expression levels in CD4 and COB T cells were measured by staining individual tags (myc and V5 respectively). Cells were treated with TMP at 1uM and GZV at 1uM. **B)** Cytotoxicity of COB+ T cells expressing the AND gate NASCAR was measured after treatment with combinations of TMP and GZV. Cells were treated with TMP at 1uM and GZV at 0.2uM (mean ± s.d., n = 3). **C)** Dose response of AND gate NASCAR-expressing CD4+ T cells when treated with increasing levels of TMP and GZV. Cell activity measured in terms of CD69, IL-2 and IFN-y (n = 3, mean values displayed).

**Figure S8:**
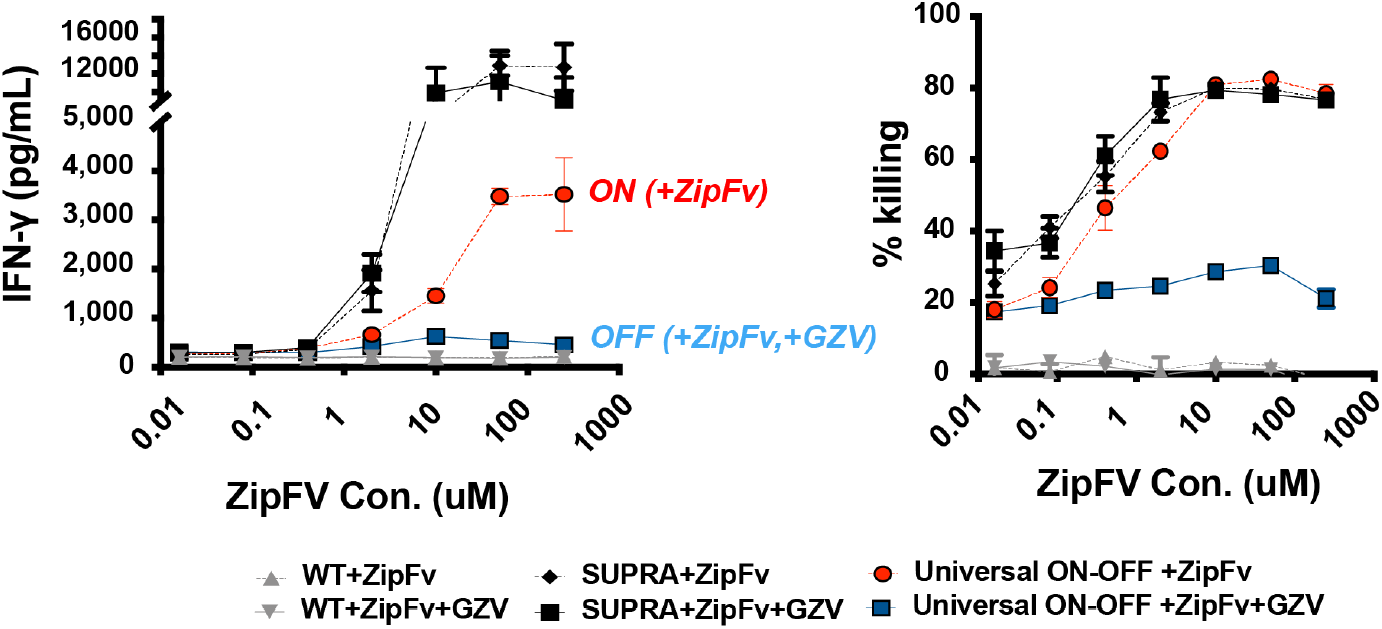
Dose response of Universal ON-OFF NASCAR to zipFv. Regulation of ON state of the SUPRA-NASCAR by different concentrations of zipFv, as mea­ sured by IFN-y (left) and cell killing ability (right). 1 uM of GZV was used for all GZV conditions to turn off SUPRA-NASCAR functionality (mean ± s.d., n = 3).

**Figure S9:**
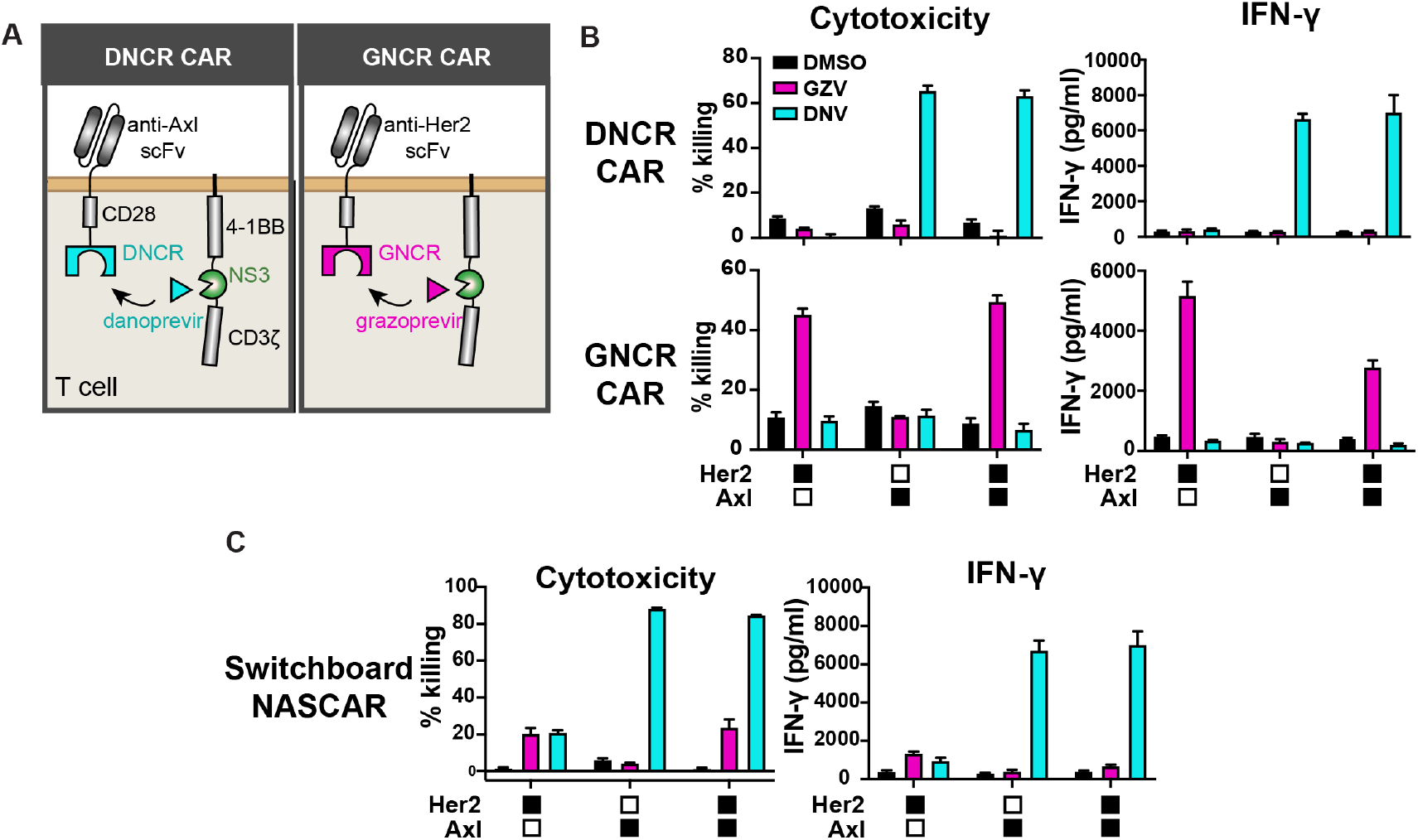
Implementation of individual and switchboard NASCARs in primary T cells. **A)** Schematic of how single NS3 reader CARs function in T cells. **B)** Cytotoxicity and cytokine levels of individual NS3 reader CARs in primary T cells when treated with various antigen-expressing target cells, and with or without NS3 inhibitor (mean ± s.d., n = 3). **C)** Cytotoxicity and cytokine levels of switchboard NASCAR in primary T cells when treated with various antigen-expressing target cells, and with or without NS3 inhibitor (mean ± s.d., n = 3)

